# DIA-Pipe: Identification and Quantification of Post-Translational Modifications using exclusively Data-Independent Acquisition

**DOI:** 10.1101/141382

**Authors:** Jesse G. Meyer, Sushanth Mukkamalla, Alexandria K. D’Souza, Alexey I. Nesvizhskii, Bradford W. Gibson, Birgit Schilling

**Affiliations:** Buck Institute for Research on Aging, Novato, CA, USA.; Department of Computational Medicine and Bioinformatics, University of Michigan, Ann Arbor, Michigan, USA.; Department of Pathology, University of Michigan, Ann Arbor, Michigan, USA.

## Abstract

Label-free quantification using data-independent acquisition (DIA) is a robust method for deep and accurate proteome quantification^1,2^. However, when lacking a pre-existing spectral library, as is often the case with studies of novel post-translational modifications (PTMs), samples are typically analyzed several times: one or more data dependent acquisitions (DDA) are used to generate a spectral library followed by DIA for quantification. This type of multi-injection analysis results in significant cost with regard to sample consumption and instrument time for each new PTM study, and may not be possible when sample amount is limiting and/or studies require a large number of biological replicates. Recently developed software (e.g. DIA-Umpire) has enabled combined peptide identification and quantification from a data-independent acquisition without any pre-existing spectral library^3,4^. Still, these tools are designed for protein level quantification. Here we demonstrate a software tool and workflow that extends DIA-Umpire to allow automated identification and quantification of PTM peptides from DIA. We accomplish this using a custom, open-source graphical user interface DIA-Pipe (https://github.com/jgmeyerucsd/PIQEDia/releases/tag/v0.1.2) (figure 1a).

---

**To the editor:** Label-free quantification using data-independent acquisition (DIA) is a robust method for deep and accurate proteome quantification^1,2^. However, when lacking a pre-existing spectral library, as is often the case with studies of novel post-translational modifications (PTMs), samples are typically analyzed several times: one or more data dependent acquisitions (DDA) are used to generate a spectral library followed by DIA for quantification. This type of multi-injection analysis results in significant cost with regard to sample consumption and instrument time for each new PTM study, and may not be possible when sample amount is limiting and/or studies require a large number of biological replicates. Recently developed software (e.g. DIA-Umpire) has enabled combined peptide identification and quantification from a data-independent acquisition without any pre-existing spectral library^3,4^. Still, these tools are designed for protein level quantification. Here we demonstrate a software tool and workflow that extends DIA-Umpire to allow automated identification and quantification of PTM peptides from DIA. We accomplish this using a custom, open-source graphical user interface DIA-Pipe (https://github.com/jgmeyerucsd/PIQEDia/releases/tag/v0.1.2) (figure 1a).

We benchmarked this all-DIA workflow using acetylated peptides enriched by immunoprecipitation (see **supplemental methods**). Strikingly, the peptide peaks identified from pseudo-MS/MS spectra are selected for integration flawlessly by Skyline^5^ according to mProphet modeling (figure 1b). This is due to the fact that in this case (unlike using external DDA libraries) targeted extraction in Skyline is performed using peptide signals already confidently detected and processed by DIA-Umpire signal extraction for pseudo-MS/MS generation, ensuring that mProphet can easily model the observed distributions. Even in rare cases where the wrong peak might be integrated by Skyline, these outliers will likely be filtered post-integration by mapDIA^6^, which employs several fragment-level interference filters. Further, the distribution of coefficients of variation (CVs) for these peptides among three technical replicates was excellent; nearly 70% of peptides had CV values below 10%, and 90% of peptides showed CVs below 20% (figure 1c). Most importantly, using mapDIA to quantify site-specific changes in acetylation resulted in very accurate and precise quantification when comparing technical replicates of 0.5X or 1X quantity injections (figure 1d). We further note that compared to the stochastic nature of identification with DDA^7^, the per-run identification numbers from DIA replicates are very consistent, i.e. 1,655 ± 29 acetylated peptides identified per injection (figure 1e).

**Figure 1:**
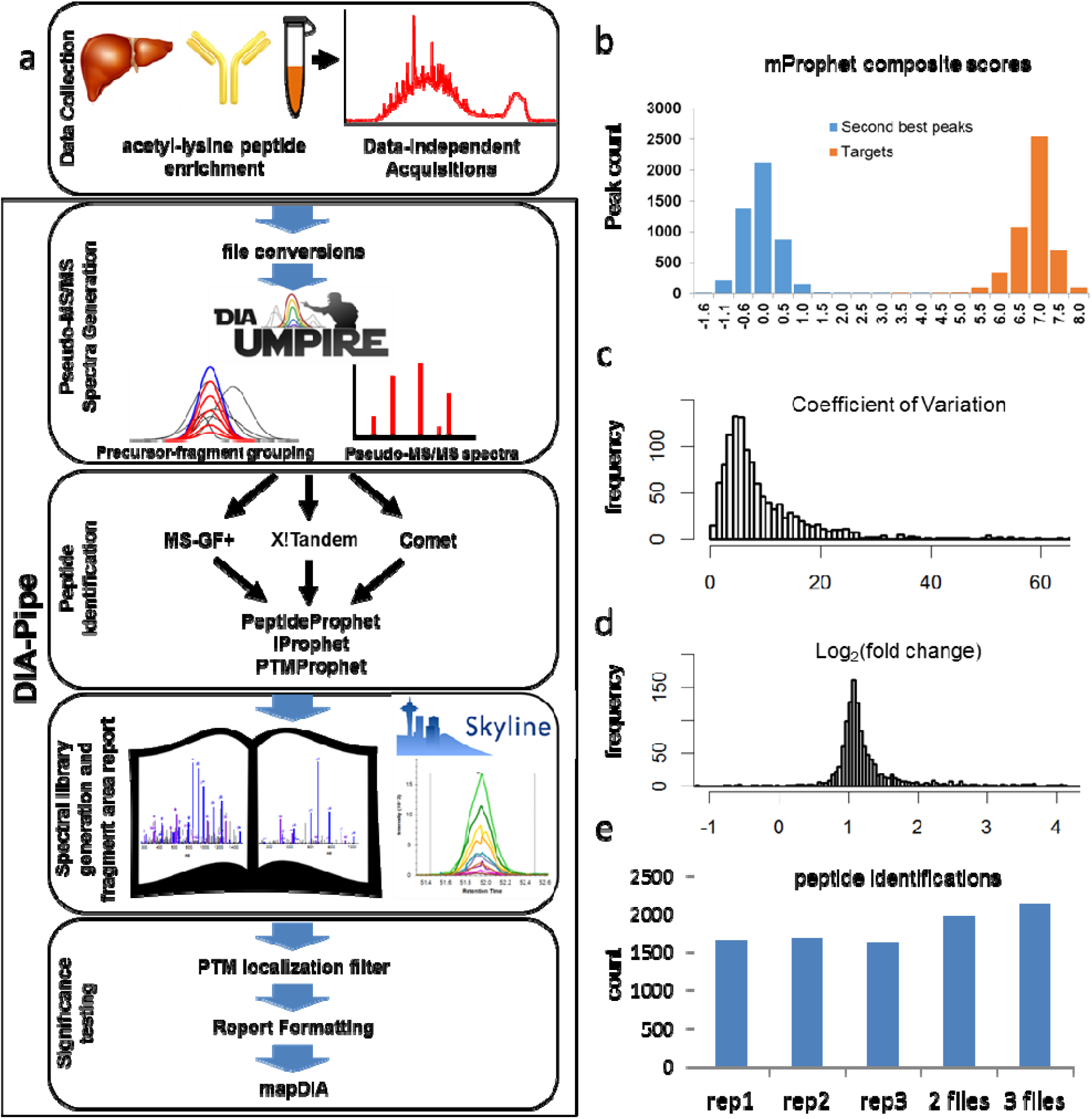
Automated Qualitative and Quantitative Analysis of Post-Translational Modifications using DIA-Pipe. **(a)** Workflow of the all-DIA strategy for identification and quantification of PTMs. Modified peptides enriched from biological samples are analyzed by data-independent acquisition. All data analysis steps starting from instrument. wiff files, acquired on a TripleTOF 5600, can be completed using the DIA-Pipe GUI, including: (1) file conversion and pseudo-MS/MS spectra generation using DIA-Umpire, (2) database searching by MS-GF+, X! Tandem, and Comet followed by results refinement and combination using PeptideProphet/iProphet/PTMProphet, (3) automated spectral library generation and fragment area extraction using SkylineRunner, and finally, (4) Skyline report filtering and formatting for significance testing with mapDIA. **(b)** mProphet composite score distributions of target and second-best peaks picked by Skylineshowing essentially error-free peak picking by Skyline of peptides identified using pseudo-MS/MS spectra. **(c)** Distribution of coefficient of variations observed from three technical replicates for 1,182 acetylation sites. **(d)** Observed distributions of log_2_(fold change) computed by mapDIA using three technical replicates of 1X injection volume compared to three technical replicates of 0.5X injection volume; expected log_2_(fold change) value =1. **(e)** Number of identified peptides from each single replicate injection and from the combination of two or three replicates. An average of 1,655 acetylated peptides were identified per replicate. The combination of two or three replicates increased the number of identifications by 19% or 29%, respectively.

Our all-DIA strategy using DIA-Pipe drastically simplifies the process of label-free PTM identification and quantification, and practically eliminates the need for subjective human data analysis and error. In addition to improved PTM quantification, because each sample is only analyzed by one DIA, total instrument acquisition time is reduced by at least twofold. Given all the benefits of this all-DIA strategy, we predominantly use the all-DIA strategy for high throughput PTM identification and label-free quantification studies.

## Acknowledgments

This work was supported by the NIDDK (R24DK085610, to B.W.G., Subcontract) and NIGNS (5R01GM094231, to A.I.N). J.G.M. was supported by an NIH T32 (T32G000266, PI: J. Campisi). We acknowledge support from the NIH shared instrumentation grant for the TripleTOF system at the Buck Institute (1S10 OD016281, to B.W.G.). The authors thank Brendan MacLean for assistance with Skyline.

## Supplemental figure 1 legend

Screenshot of the DIA-Pipe GUI. The software has several optional modules that can be run, where each subsequent module can automatically use the output from the previous module. The leftmost section is used to specify parameters for the DIA-Umpire signal extraction module that generates pseudo-MS/MS spectra for searching. The middle section is used to set the database search parameters for up to three database searches. The right section contains input for PeptideProphet, iProphet, PTMProphet, Skyline peak area report generation, and mapDIA significance testing. Additionally, the user can save and load up to four default parameter sets using the save/load buttons.

